# SARS-CoV-2 spike protein induces the cytokine release syndrome by stimulating T cells to produce more IL-2

**DOI:** 10.1101/2023.11.01.565098

**Authors:** Chao Niu, Tingting Liang, Yongchong Chen, Shan Zhu, Lei Zhou, Naifei Chen, Lei Qian, Yufeng Wang, Min Li, Xin Zhou, Jiuwei Cui

## Abstract

Cytokine release syndrome (CRS) is one of the leading causes of mortality in COVID-19 patients caused by the SARS-CoV-2 coronavirus. However, the mechanism of CRS induced by SARS-CoV-2 is vague. This study shows that dendritic cells loaded with spike protein of SARS-CoV-2 stimulate T cells to release much more IL-2, which subsequently cooperates with spike protein to facilitate peripheral blood mononuclear cells to release IL-1β, IL-6, and IL-8. These effects are achieved via IL-2 stimulation of NK cells to release TNF-α and IFN-γ, as well as T cells to release IFN-γ. Mechanistically, IFN-γ and TNF-α enhance the transcription of CD40, and the interaction of CD40 and its ligand stabilizes the membrane expression of TLR4 which serves as a receptor of spike protein on the surface of monocytes. As a result, there is a constant interaction between spike protein and TLR4, leading to continuous activation of NF-κB. Furthermore, TNF-α also activates NF-κB signaling in monocytes, which further cooperates with IFN-γ and spike protein to modulate NF-κB-dependent transcription of CRS-related inflammatory cytokines. Targeting TNF-α/IFN-γ in combination with TLR4 may represent a promising therapeutic approach for alleviating CRS in individuals with COVID-19.

## Introduction

COVID-19, caused by SARS-CoV-2, has become a global pandemic since its outbreak in 2019.(Hu et al., 2021) As of 12 April 2023, there have been 762,791,152 confirmed cases of COVID-19, including 6,897,025 deaths, reported to WHO. The clinical manifestations of severe COVID-19 are diverse, with acute respiratory distress syndrome, cytokine release syndrome (CRS), multiple organ failure, and death being the most notable.(Delorey et al., 2021; Wang & Perlman, 2022) CRS-related cytokines tend to increase progressively with the severity of the disease (Xiao et al., 2021) and may be the leading cause of life-threatening respiratory diseases in severe COVID-19 patients.(Que et al., 2022) Interestingly, it has been observed that the concentration of CRS-related cytokines is higher in the plasma of COVID-19 patients admitted to the intensive care unit (ICU) than in those who are not.(Huang et al., 2020) These cytokines include tumor necrosis factor (TNF)-α, interleukin (IL)-1β, IL-6, IL-10, IL-17, interferon (IFN)-γ and IL-2, etc.(Giamarellos-Bourboulis et al., 2020; Mehta et al., 2020) However, the interplay between these cytokines remains unclear. Understanding the relationship between various cytokines in CRS is crucial for developing targeted therapies for COVID-19 and other cytokine storm syndromes.

Spike protein is one of the four main structural proteins of SARS-CoV-2, (Sen et al., 2021) which plays a crucial role in the virus’s ability to enter the host cells as it can bind to the ACE2 receptor.(Jackson et al., 2022) It also binds directly to pattern recognition receptor TLR4 to activate downstream signaling pathways which upregulate inflammatory factors such as IL-1β and IL-6.(Zhao et al., 2021a, 2021b) Therefore, spike protein may be a critical factor contributing to CRS in COVID-19 patients. The amount of spike protein increases with the increase of viral load. However, clinical studies have shown no significant difference in viral load between severe and mild COVID-19 patients, (To et al., 2020) and the viral load of patients with COVID-19 is relatively high in the initial stage of infection.(Pan et al., 2020; Walsh et al., 2020) These studies show that the accumulation of viral antigens may not be the leading cause of CRS. These findings indicate that factors other than the viral load may also significantly induce CRS in COVID-19 patients. The intricate interplay between spike protein, immune cells, and cytokines during the CRS remains largely unknown, highlighting the urgent need for further investigation.

During SARS-CoV-2 infection, alveolar macrophages play a critical role in detecting the virus and producing cytokines and chemokines to recruit innate and adaptive immune cells for virus elimination and disease prevention.(Gajjela & Zhou, 2022) Of note, macrophages and certain monocyte subsets are believed to be the decisive cells of CRS in patients with severe COVID-19.(Liao et al., 2020) Moreover, the over-activation of inflammatory response by other immune cells such as neutrophils, Dendritic cells (DCs), natural killer (NK) cells, B cells, and T cells can also contribute to CRS in this context.(Tan & Tang, 2021) Nonetheless, further research is still needed to determine the precise involvement of these cell types in CRS.

In this study, we investigated the mechanism underlying the development of CRS induced by SARS-Cov-2. Our finding suggests a cooperative effect of IL-2 and spike protein in stimulating peripheral blood mononuclear cells (PBMCs) to secrete IL-1β, IL-6, and IL-8. Mechanistically, DCs loaded with spike protein stimulate T cells to secrete IL-2, which subsequently facilitates the production of TNF-α and IFN-γ by NK cells and IFN-γ by T cells. Together, TNF-α and IFN-γ make monocytes more active, and when stimulated by spike protein of SARS-CoV-2, these cells can release more CRS-related cytokines. Overall, our findings provide insights into the possible mechanism underlying CRS in COVID-19 patients and illustrate the complex interplay among various cytokines. These findings may pave the way for developing novel therapeutic targets for treating CRS, thereby offering promising avenues for clinical interventions in patients with COVID-19.

## Results

### IL-2 cooperates with spike protein to stimulate PBMCs to secrete IL-1β, IL-6, and IL-8

In order to analyze whether spike protein can stimulate PBMCs to secrete IL-6 or IL-1β, PBMCs were treated with different concentrations of spike protein, and a dose-dependent increase in the secretion of IL-6 and IL-1β was observed (Figure 1A). Hereafter, 10 nM spike protein was used for further experiments unless otherwise specified. The stimulatory effect of spike protein was further confirmed by increased secretion of IL-1β, IL-6, and IL-8 upon treatment with spike protein in PBMCs (Figure S1). However, no significant changes were observed in the expressions of other cytokines such as IL-2, IL-4, IL-5, IL-10, IL-12, IL-17, TNF-α, IFN-α, or IFN-γ (Figure S1). These results suggest that spike protein alone cannot stimulate PBMCs to release various inflammatory factors. This phenomenon differs from the induction of diverse cytokine release in severe patients with COVID-19, indicating that other antigens of COVID-19 or other immune cells may be involved in the process of CRS in the human body.

**Figure 1.**
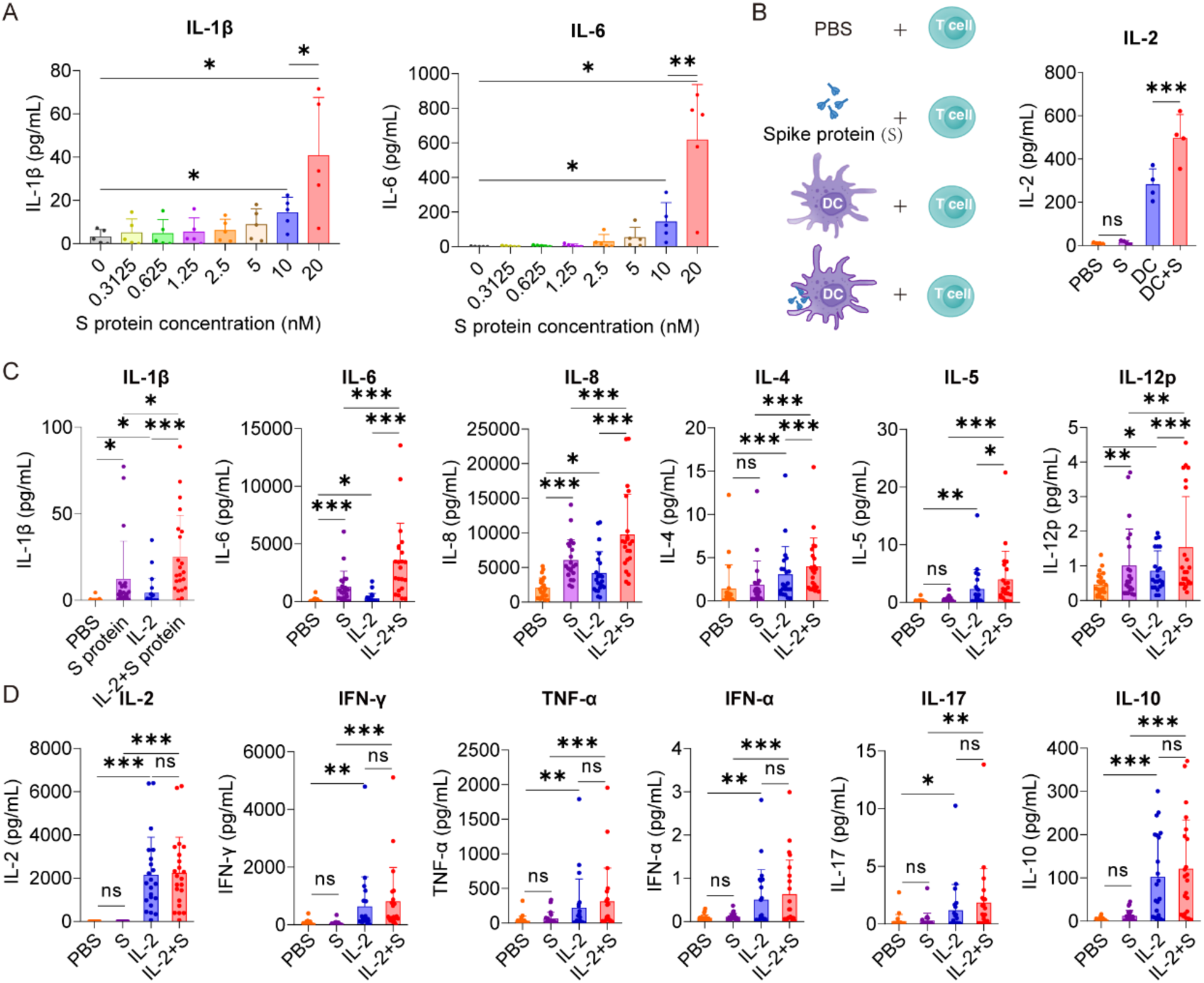
IL-2 cooperates with spike protein to stimulate PBMCs to secrete IL-6, IL-1β, and IL-8. (A) Quantifying concentrations of IL-1β and IL-6 in supernatants of PBMCs stimulated by different concentrations of spike protein for 16 hrs according to CBA (n = 5 biological replicates). (B) Bar graph (right) showing the concentration of IL-2 in the supernatants of T cells cocultured with DC, DC activated with spike protein, spike protein, or PBS as indicated by the schema (left). n = 4 biological replicates. (C, D) Quantifying concentrations of cytokines by CBA in the supernatants of PBMCs treated with PBS, spike protein, IL-2, or IL-2 combined with spike protein for 16 hrs (n = 22 biological replicates). Data are presented as mean ± SD. ns: not significant, *p < 0.05, **p < 0.01, and ***p < 0.001 as analyzed by one-way ANOVA.

Among various cytokines involved in the CRS of COVID-19 patients, IL-2 has drawn our particular attention. This is due to the noticeable elevation of IL-2 in the plasma of many severe COVID-19 patients (Akbari et al., 2020; Huang et al., 2020; J. Liu et al., 2020) and high-dose IL-2 administration causes capillary leak syndrome, (Boyman & Sprent, 2012; van Haelst Pisani et al., 1991) a severe clinical manifestations of CRS.(Case et al., 2020) Although the source of IL-2 remains elusive, these studies hint at the crucial role of IL-2 in developing CRS in severe COVID-19 patients. Spike protein is a virus antigen which can be recognized by antigen-producing cells, such as DCs. We examined the effect of DCs loaded with spike protein on T-cell activation and found that DCs loaded with spike protein can stimulate T cells to secrete higher levels of IL-2 than did control DCs (Figure 1B). Along this line, we observed an increased secretion of IL-1β, IL-6, IL-8, IL-4, IL-5, and IL-12 from PBMCs treated with the combination of IL-2 and spike protein, as compared with spike protein or IL-2 alone (Figure 1C). In contrast, IL-2 alone stimulated PBMCs to secrete IFN-γ, TNF-α, IFN-α, IL-17, and IL-10, while the combination of spike protein and IL-2 did not show any synergistic effect on the secretion of these cytokines (Figure 1D). Together, these data suggest that IL-2 released by T cells activated by DCs stimulated with spike protein may serve as an amplifier in inducing CRS in COVID-19 patients in a manner of cooperation with spike protein.

### Monocytes are the critical cells for the cooperation between IL-2 and spike protein in stimulating PBMCs to secrete inflammatory cytokines

IL-6 is a critical cytokine involved in the pathogenesis of CRS in COVID-19 patients. In severe cases of COVID-19, elevated levels of IL-6 have been observed and are believed to contribute to the systemic inflammatory response and organ damage seen in some patients.(B. Liu et al., 2020; Smoke et al., 2021) Therefore as a readout, IL-6 was used to analyze which cells in PBMCs are responders or effectors of the synergistic effect of IL-2 and spike protein to release cytokines. Intracellular staining and flow cytometry analysis revealed that spike protein, rather than IL-2, significantly increased the production of IL-6 in monocytes in PBMCs, and the combination of spike protein and IL-2 further enhanced this effect (Figure 2A). However, no increase in the expression level of IL-6 was observed in NK cells, B cells, or T cells (Figure 2B). These findings indicate that IL-2 can stimulate monocytes in PBMCs to produce more IL-6 in cooperation with spike protein. To further determine the involvement of monocytes, they were removed from PBMCs and then stimulated with spike protein, IL-2, or spike protein combined with IL-2. Compared with in that PBMCs, the levels of various cytokines, including IL-1β, IL-6, IL-8, IL-4, IL-5, IL-12, IL-2, TNF-α, and IL-17 decreased significantly in the supernatant of PBMCs without monocytes after treatment with spike protein combined with IL-2. Of note, IL-1β, IL-6, and IL-8 levels also decreased significantly in the supernatant of PBMCs treated with spike protein after monocytes removal (Figure 2C). These results indicate that monocytes play an essential role in the cytokine secretion by PBMCs stimulated by spike protein or IL-2 in cooperation with spike protein.

**Figure 2.**
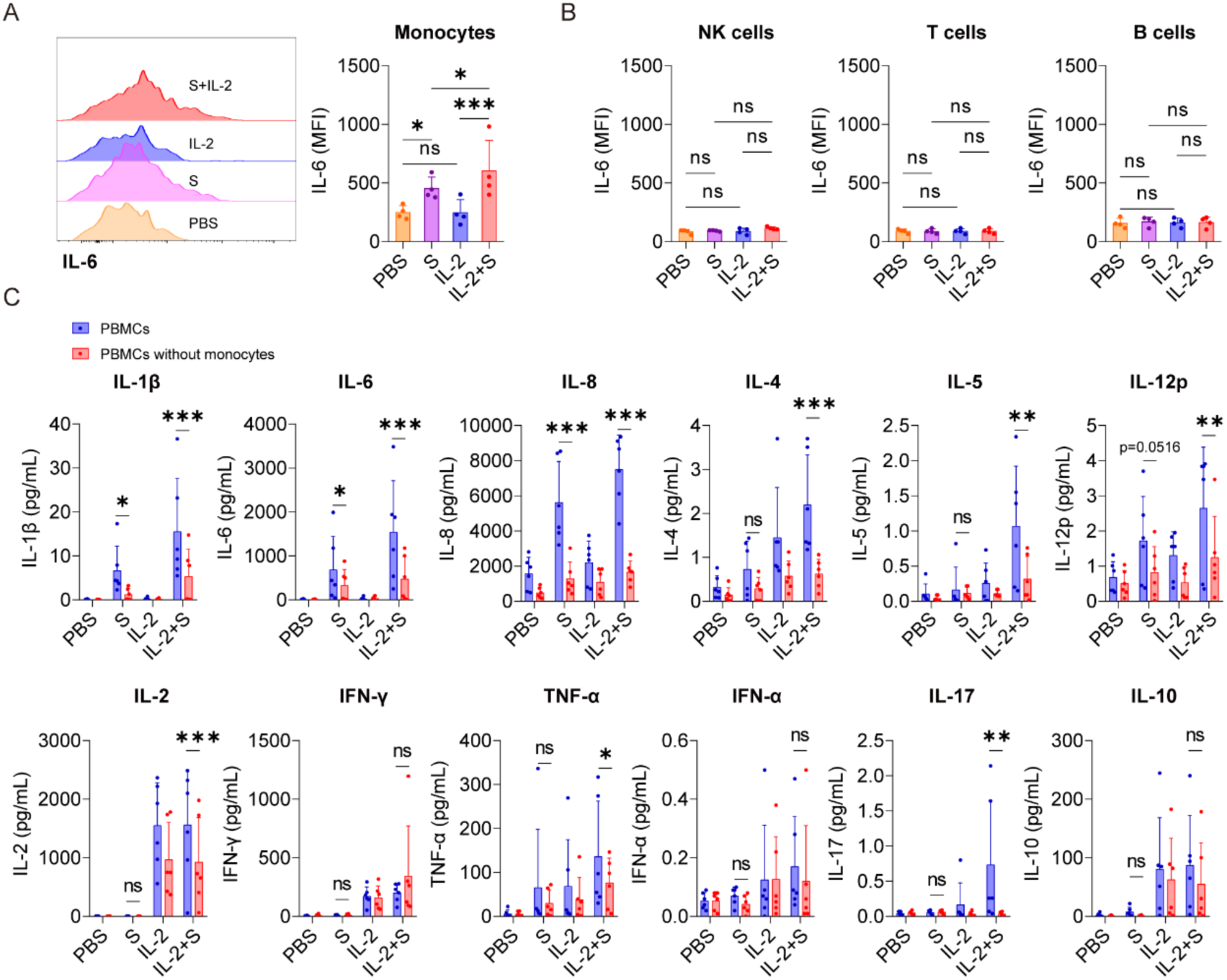
Monocytes are pivotal cells in the secretion of CRS-related cytokines in PBMCs co-stimulated with IL-2 and spike protein. (A) Representative intracellular staining analysis (left) and quantification (right) of the expression of IL-6 by flow cytometry in monocytes of PBMCs stimulated with PBS, spike protein, IL-2, or spike protein combined with IL-2 for 16 hrs (n = 4 biological replicates). MFI, mean fluorescence intensity. (B) Quantification of the expression of IL-6 in NK cells, T cells, and B cells of PBMCs described in A. (C) Quantifying the concentrations of cytokines by CBA in the supernatants of PBMCs and PBMCs without monocytes stimulated with PBS, spike protein, IL-2, or spike protein combined with IL-2 for 16 hrs (n = 6 biological replicates). Data are presented as mean ± SD. ns: not significant, *p < 0.05, **p < 0.01, and ***p < 0.001 as analyzed by one-way ANOVA (A, B) or two-way ANOVA (C).

### IL-2 activates NF-κB of monocytes via stimulating PBMCs to release TNF-α and IFN-γ

To investigate the mechanism underlying the synergistic effect of IL-2 and spike protein in inducing CRS, transcriptomic analysis was conducted on PBMCs treated with PBS, spike protein, IL-2, or a combination of spike protein and IL-2. Our analysis revealed that stimulation with spike protein activated multiple signaling pathways, including NF-κB, TNF, and JAK-STAT in PBMCs (Figure S2A-C). Notably, the combination of IL-2 and spike protein further amplified the activation of NF-κB, TNF, Toll-like receptor, and JAK-STAT signaling pathways compared to spike protein alone (Figure 3A-C). Flow cytometry analysis showed that spike protein facilitated the phosphorylation of p65, an indicator of activation of the NF-κB signaling pathway, in monocytes within PBMCs, and IL-2 combined with spike protein further enhanced this effect (Figure 3D and S2D).

**Figure 3.**
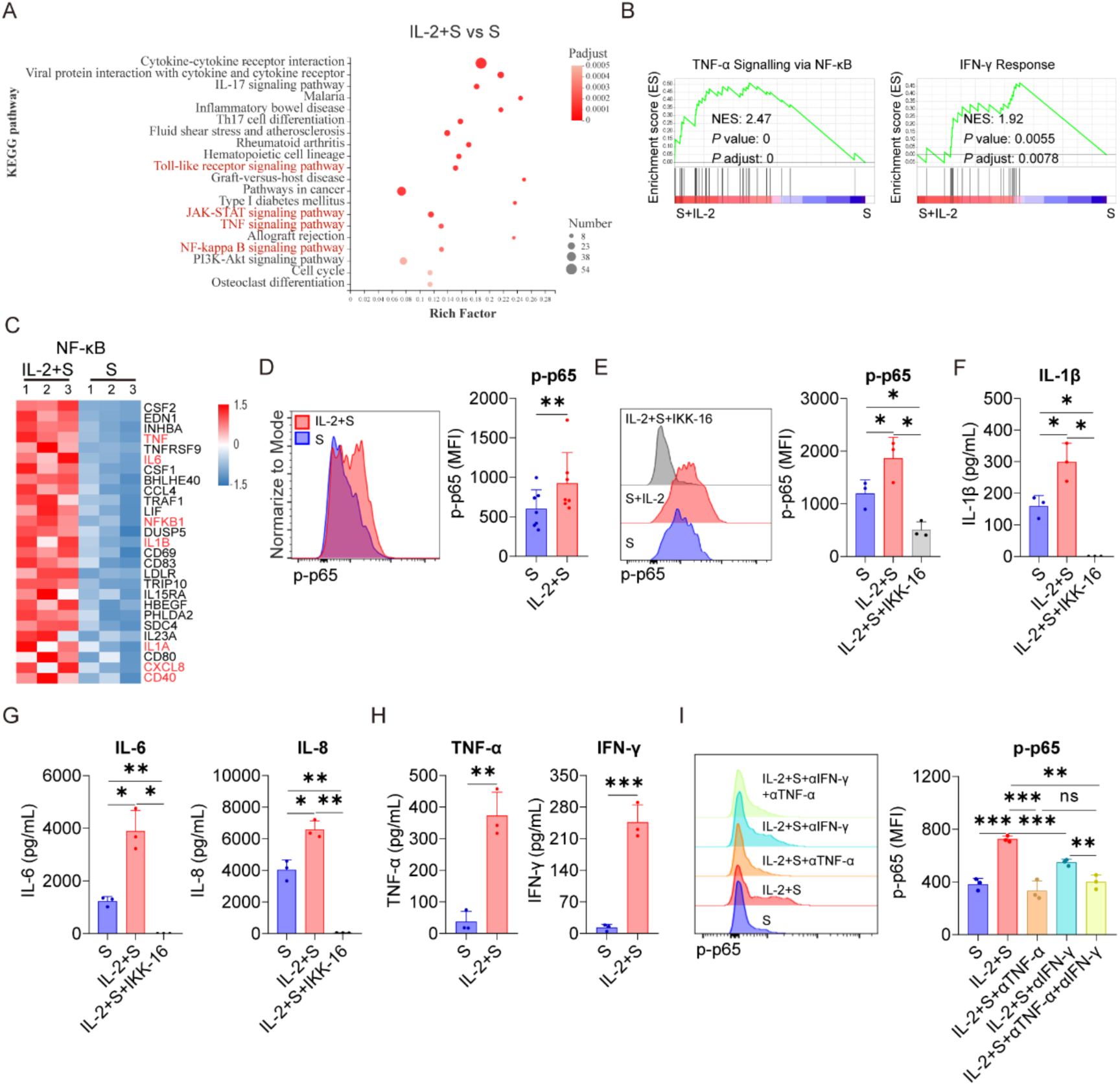
TNF-α and IFN-γ mediate the activation of NF-κB in PBMCs upon the synergistic stimulation by spike protein and IL-2. (A) Bubble plot showing the enrichment of KEGG pathways based on the transcriptomic analysis of PBMCs treated with IL-2 combined with spike protein or spike protein alone for 16 hrs (n = 3 biological replicates). (B) GSEA shows the TNF-α signaling via NF-κB and the IFN-γ signatures in the transcriptomic analysis described in A. NES: normalized enrichment score; FDR: false discovery rate. (C) Heat map showing the differentially expressed genes in the NF-κB pathway in the samples in A. (D) Representative intracellular staining analysis (left) and quantification (right) of the expression of p-p65 by flow cytometry in monocytes of PBMCs stimulated with spike protein or spike protein combined with IL-2 for 16 hrs (n = 7 biological replicates). (E) Representative intracellular staining analysis (left) and quantification (right) of the expression of p-p65 in monocytes of PBMCs stimulated with spike protein, spike protein combined with IL-2, or spike protein combined with IL-2 and IKK-16 for 16 hrs (n = 3 biological replicates). (F, G) Quantifying concentrations of IL-1β, IL-6, and IL-8 by CBA in the supernatants of PBMCs in E. (H) Quantifying concentrations of TNF-α and IFN-γ in the supernatants of PBMCs stimulated with spike protein or spike protein combined with IL-2 for 16 hrs (n = 3 biological replicates). (I) Representative intracellular staining analysis (left) and quantification (right) of the expression of p-p65 in monocytes of PBMCs stimulated with spike protein, spike protein combined with IL-2 and control IgG, or spike protein combined with IL-2 and TNF-α or/and IFN-γ blocking antibodies for 16 hrs (n = 3 biological replicates). Data are presented as mean ± SD. ns: not significant, *p < 0.05, **p < 0.01, and ***p < 0.001 as analyzed by paired Student’s t-test (D and H) or one-way ANOVA (E, G, and I).

Significantly, inhibiting the activation of NF-κB with IKK-16 not only significantly reduced the secretion of IL-6, IL-1β, and IL-8 by PBMCs stimulated by spike protein (Figure S2E, S2F) but also blocked the secretion of these cytokines by PBMCs stimulated by IL-2 in cooperation with spike protein (Figure 3E-G). As TNF and JAK-STAT signaling pathways were significantly enriched in PBMCs treated with IL-2 combined with spike protein compared to spike protein alone (Figure 3A, 3B), it appears that IL-2 may regulate these pathways in PBMCs. The JAK-STAT signaling pathway is tightly regulated by IFN and plays an important role in various biological processes.(Di Bona et al., 2006; Li et al., 2023; O’Connell et al., 2015) Interestingly, in the IL-2 combined with spike protein group, TNF-α and IFN-γ levels were significantly increased compared to the spike protein group (Figure 3H). IL-2 alone elevated TNF-α and IFN-γ levels in PBMCs, too (Figure 1D). Importantly, blocking TNF-α or/and IFN-γ with antibodies during PBMCs stimulation with IL-2 combined with spike protein inhibited the activation of the NF-κB signaling pathway (Figure 3I).

Based on these findings, it can be inferred that IL-2 plays a crucial role in activating the NF-κB signaling pathway in PBMCs by promoting the secretion of TNF-α and IFN-γ. Previous studies have reported that spike protein also activates the NF-κB signaling pathway.(Forsyth et al., 2022) Therefore, the interplay between IL-2 and TNF-α/IFN-γ might be crucial in mediating the synergistic effect of IL-2 and spike protein in inducing CRS in COVID-19 patients.

### TNF-α and IFN-γ cooperate with spike protein to stimulate monocytes to secrete IL-1β, IL-6, and IL-8

As a critical transcription factor, NF-κB is involved in producing of many inflammatory cytokines, such as IL-1β and IL-6.(Al-Griw et al., 2022) Since the activation of NF-κB in PBMCs induced by IL-2 relies on TNF-α and IFN-γ (Figure 3I), we sought to explore whether the secretion of CRS-related cytokines by PBMCs stimulated with IL-2 and spike protein also depends on these cytokines. Our results demonstrated that the TNF-α blocking antibodies effectively suppressed the secretion of IL-1β, IL-6, and IL-8 in PBMCs stimulated by IL-2 and spike protein. Similarly, the IFN-γ blocking antibodies inhibited the secretion of IL-1β and IL-6 in PBMCs stimulated under the same condition. Interestingly, the combined blocking effect of TNF-α and IFN-γ was not significantly different from that of single blocking antibodies for TNF-α (Figure 4A), suggesting that TNF-α plays a more important role in mediating IL-2-induced cytokines secretion, especially IL-8. Moreover, TNF-α or IFN-γ, when combined with spike protein, stimulated the secretion of CRS-related cytokines in PBMCs, with the effect of their combination being the most potent (Figure 4B). Finally, we observed that IKK-16, an NF-κB inhibitor, completely inhibited the secretion of IL-1β, IL-6, and IL-8 by PBMCs stimulated with TNF-α or IFN-γ that is cooperating with spike protein (Figure S3). Collectively, TNF-α and IFN-γ are indispensable in activating NF-κB and inducing CRS-related cytokines in PBMCs via cooperating with spike protein.

**Figure 4.**
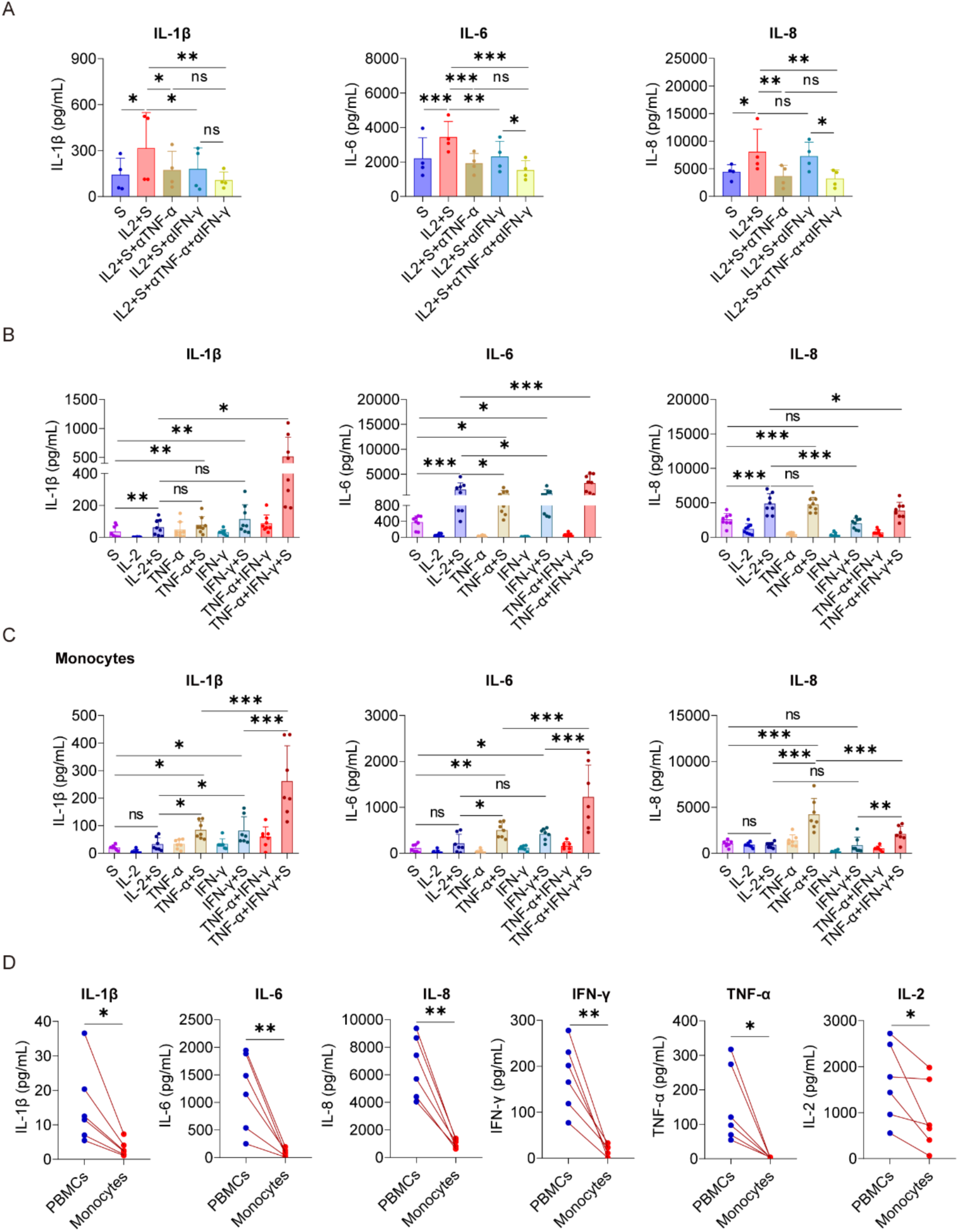
TNF-α and IFN-γ cooperate with spike protein to stimulate monocytes to release IL-1β, IL-6, and IL-8. (A) Quantifying concentrations of IL-1β, IL-6, and IL-8 by CBA in the supernatants of PBMCs stimulated with spike protein, spike protein combined with IL-2 together with/without blocking antibodies against TNF-α or IFN-γ for 16 hrs (n = 4 biological replicates). (B) Quantifying concentrations of IL-1β, IL-6, and IL-8 by ELISA in the supernatants of PBMCs stimulated with spike protein, IL-2, TNF-α, IFN-γ, or spike protein combined with these cytokines for 16 hrs (n = 8 biological replicates). (C) Quantifying concentrations of IL-1β, IL-6, and IL-8 by CBA in the supernatants of monocytes stimulated with spike protein, IL-2, TNF-α, IFN-γ, or spike protein combined with these cytokines for 16 hrs. (n = 7 biological replicates). (D) Quantifying concentrations of IL-1β, IL-6, IL-8, IFN-γ, TNF-α, and IL-2 by CBA in the supernatants of PBMCs and monocytes stimulated with spike protein combined with IL-2 for 16 hrs. Each red line connects PBMCs and monocytes from the same donor. (n = 6 biological replicates) Data are presented as mean ± SD. ns: not significant, *p < 0.05, **p < 0.01, and ***p < 0.001 as analyzed by one-way ANOVA (A, B, and C) or paired Student’s t-test (D).

As we demonstrated that monocytes are the critical cells for the cooperation between IL-2 and spike protein in stimulating PBMCs to secrete inflammatory cytokines (Figure 2), we further tested the synergistic effect of TNF-α/IFN-γ and spike protein in purified monocytes. As expected, TNF-α or IFN-γ, when combined with spike protein, stimulated monocytes to secrete IL-1β, IL-6, and IL-8, with the most potent effect observed when spike protein was combined with TNF-α and IFN-γ (Figure 4C). However, surprisingly, there was no synergism observed when IL-2 was combined with spike protein in purified monocytes (Figure 4C). These observations suggest that other immune cells are involved in mediating the synergistic effect of IL-2 and spike protein in PBMCs. Therefore, to test this hypothesis, we treated PBMCs and monocytes from the same volunteer with IL-2 and spike protein, and then measured cytokine levels in the supernatant. The results showed that the expressions of IL-1β, IL-6, and L-8 in the supernatant of the monocyte group were remarkably lower than those in the PBMCs group from the same volunteer (Figure 4D). Of note, the induction of TNF-α and IFN-γ was also strikingly lower in monocytes than in the PBMCs under the same stimulatory conditions, raising the possibility that the less potent effect of IL-2 and spike protein in monocytes as compared to PBMCs is due to the lower induction of TNF-α and IFN-γ (Figure 4C). Overall, these results indicate that IL-2 and spike protein work together to stimulate other immune cells to release TNF-α and IFN-γ, thereby facilitating monocytes to secrete IL-1β, IL-6, and IL-8.

### NK cells and T cells play essential roles in the synergistic stimulation of CRS-related cytokines by IL-2 and spike protein via IFN-γ and TNF-α

To identify the immune cells responsible for secreting TNF-α and IFN-γ in PBMCs stimulated by IL-2, we conducted intracellular staining to analyze the secretion of these cytokines in NK cells, T cells, B cells, and monocytes within PBMCs. Our results indicated that compared with the PBS control group, IL-2 could increase the secretion of TNF-α in NK cells and IFN-γ in NK cells and T cells. Compared with the spike protein group, IL-2 combined with spike protein can promote the secretion of TNF-α in NK cells and monocytes and IFN-γ in NK cells and T cells (Figure 5A, B). To further investigate the role of NK cells and T cells in the secretion of inflammatory factors by PBMCs stimulated by IL-2 and spike protein, we cocultured monocytes with NK cells or T cells, respectively, and then stimulated them with IL-2 and spike protein. Our finding showed that combining IL-2 and spike protein significantly elevated IL-1β, IL-6, and IL-8 in monocytes cocultured with T cells and those cocultured with NK cells (Figure 5C, D). However, stimulating monocytes alone did not have the same effect (Figure 5C). These data further support the notion that IL-2, in collaboration with spike protein, stimulates PBMCs to secrete IL-1β, IL-6, and IL-8, and this is dependent on the presence of NK cells and T cells.

**Figure 5.**
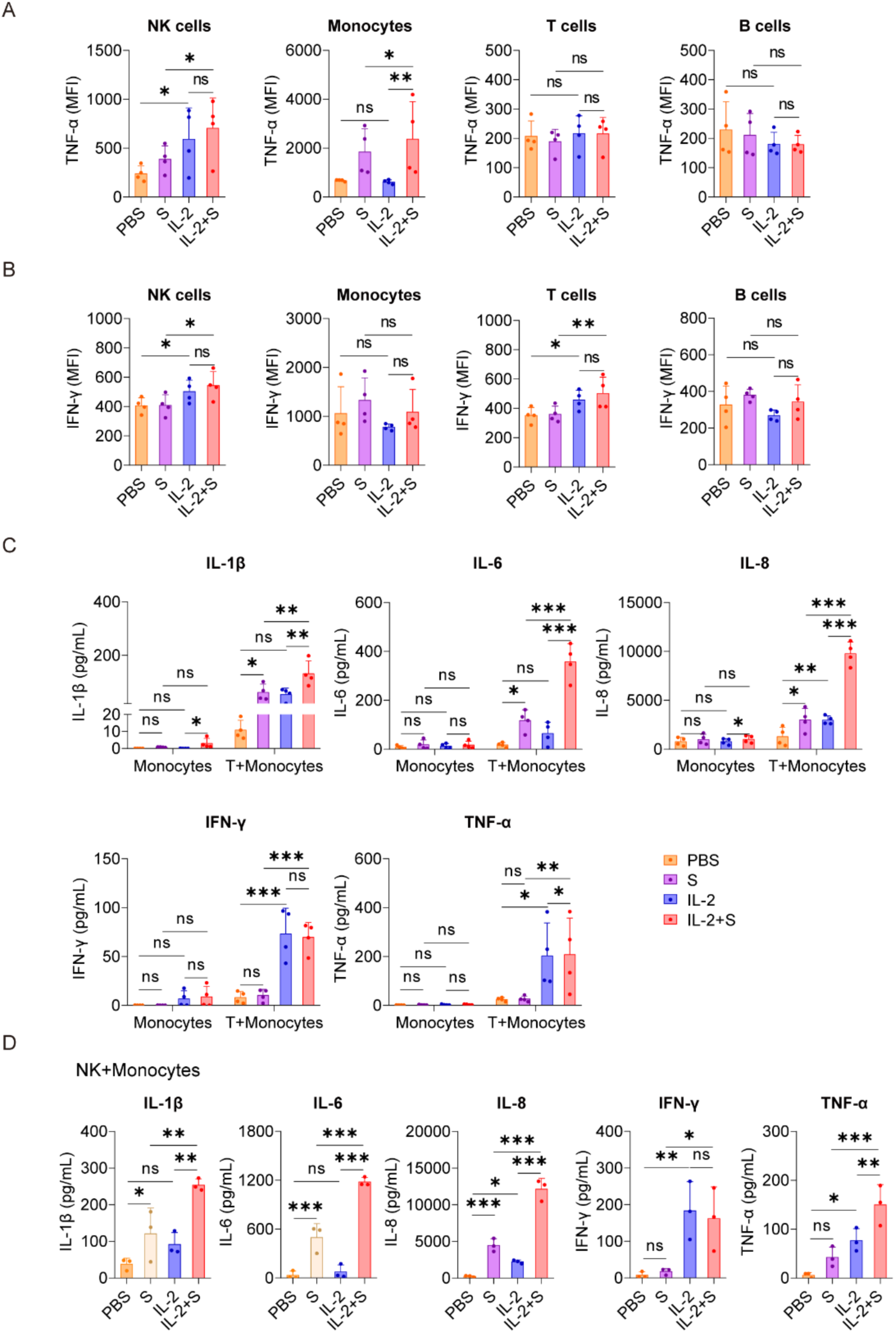
T cells and NK cells play essential roles in the synergistic stimulation of CRS-related cytokines by IL-2 and spike protein via IFN-γ and TNF-α. (A, B) Quantification of the expression of TNF-α and IFN-γ by flow cytometry in NK cells, monocytes, T cells, and B cells of PBMCs stimulated with PBS, spike protein, IL-2, or spike protein combined with IL-2 for 16 hrs (n = 4 biological replicates). (C) Quantifying concentrations of IL-1β, IL-6, IL-8, IFN-γ, and TNF-α by CBA in the supernatants of monocytes with/without cocultivation of T cells from the respective donor under stimulation with PBS, spike protein, IL-2, or spike protein combined with IL-2 for 16 hrs (n = 4 biological replicates). (D) Quantifying concentrations of IL-1β, IL-6, IL-8, IFN-γ, and TNF-α by CBA in the supernatants of monocytes cocultured with NK cells from the respective donor under stimulation with PBS, spike protein, IL-2, or spike protein combined with IL-2 for 16 hrs (n = 3 biological replicates). Data are presented as mean ± SD. ns: not significant, *p < 0.05, **p < 0.01, and ***p < 0.001 as analyzed by one-way ANOVA.

### IL-2 induce an increase in the expression of CD40 and in turn facilitates the surface localization of TLR4 in monocytes via TNF-α and IFN-γ

As we have demonstrated that NF-κB plays a critical role in mediating the synergistic effect of spike protein with IL-2, TNF-α, or IFN-γ (Figure 3E, F and S3), we sought to explore its upstream responder to spike protein and these cytokines. Through high-throughput sequencing technology, we discovered significant activation of the Toll-like receptor signaling pathway in PBMCs in the present IL-2 combined with spike protein when compared to spike protein alone (Figure 3A, 6A). It has been reported that TLR4 is a receptor of spike protein (Zhao et al., 2021a, 2021b). As IL-1β and IL-6 are classic downstream of the TLR4-NF-κB signaling cascade, IL-2, TNF-α, and IFN-γ may exert their effects via this pathway as well. We found that spike protein reduces the membrane surface expression of TLR4 in monocytes of PBMCs (Figure 6B) without affecting its transcription (Figure 6C). Spike protein binding to TLR4 leads to the internalization of TLR4, which is consistent with the effect of LPS on TLR4. Interestingly, IL-2 combined with spike protein was found to reverse the effect of spike protein on decreasing TLR4 membrane expression without affecting its transcription (Figure 6B, C).

**Figure 6.**
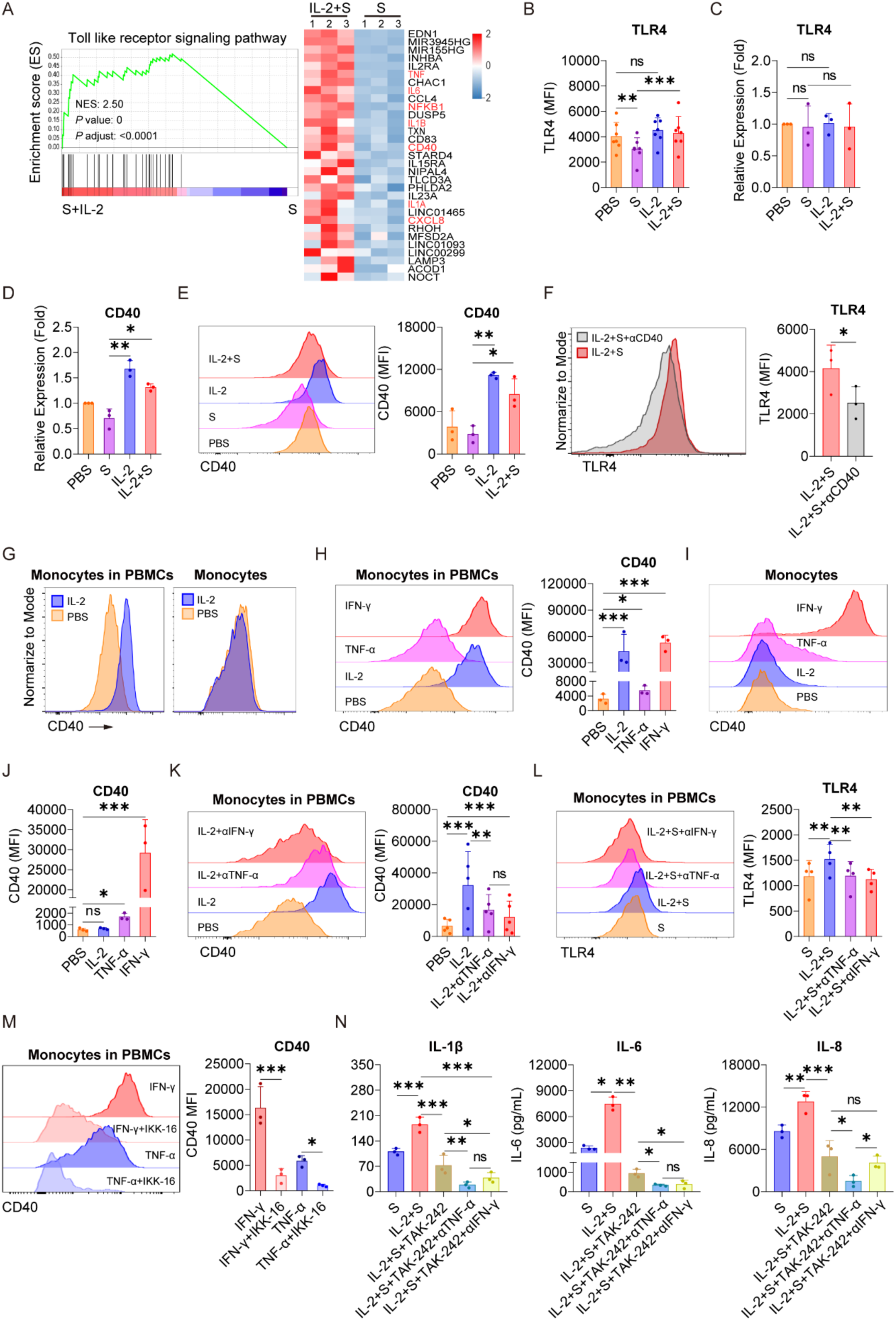
IL-2 induces an increase in the expression of CD40 and in turn facilitates the surface localization of TLR4 in monocytes via TNF-α and IFN-γ. (A) GSEA and heat map showing the differentially expressed genes of TLR signaling pathway based on transcriptomic analysis in PBMCs treated with spike protein combined with IL-2 and spike protein alone for 16 hrs (n = 3 biological replicates). (B) Quantification of the expression of TLR4 by flow cytometry on monocytes in PBMCs treated with PBS, spike protein, IL-2, or IL-2 combined with spike protein for 16 hrs (n = 7 biological replicates). (C, D) Quantification of the expression of TLR4 and CD40 by qPCR analysis in purified monocytes treated with PBS, spike protein, IL-2, or IL-2 combined with spike protein for 16 hrs (n = 3 biological replicates). (E) Representative staining analysis (left) and quantification (right) of the expression of CD40 by flow cytometry on monocytes in PBMCs stimulated with PBS, spike protein, IL-2, or spike protein combined with IL-2 for 16 hrs (n = 3 biological replicates). (F) Representative staining analysis (left) and quantification (right) of the expression of TLR4 on monocytes of PBMCs stimulated with spike protein combined with IL-2, along with the addition of CD40 blocking antibody or control IgG (n = 3 biological replicates). (G) Representative staining analysis of the expression of CD40 by flow cytometry on monocytes in PBMCs (left) and purified monocytes (right) treated with PBS or IL-2 for 16 hrs. (H) Representative staining analysis (left) and quantification (right) of the expression of CD40 by flow cytometry on monocytes in PBMCs stimulated by IL-2, TNF-α, or IFN-γ for 16 hrs (n = 3 biological replicates). (I, J) Representative staining analysis (left) and quantification (right) of the expression of CD40 by flow cytometry on purified monocytes stimulated by IL-2, TNF-α, or IFN-γ for 16 hrs (n = 3 biological replicates). (K) Representative staining analysis (left) and quantification (right) of the expression of CD40 by flow cytometry on monocytes in PBMCs stimulated with IL-2 along with the addition of blocking antibodies against TNF-α or IFN-γ or control IgG for 16 hrs (n = 5 biological replicates). (L) Representative staining analysis (left) and quantification (right) of the expression of TLR4 by flow cytometry on monocytes in PBMCs stimulated with IL-2 combined with spike protein, along with the addition of blocking antibodies against TNF-α or IFN-γ or control IgG for 16 hrs (n = 4 biological replicates). (M) Representative staining analysis (left) and quantification (right) of the expression of CD40 by flow cytometry on monocytes in PBMCs stimulated with TNF-α or IFN-γ along with the addition of NF-kB inhibitor IKK-16 for 16 hrs (n = 3 biological replicates). (N) Quantifying concentrations of IL-1β, IL-6, and IL-8 by CBA in the supernatants of PBMCs stimulated with spike protein combined with IL-2, along with the addition of TLR4 inhibitor and blocking antibodies against TNF-α or IFN-γ or control IgG for 16 hrs (n = 3 biological replicates). Data are presented as mean ± SD. ns: not significant, *p < 0.05, **p < 0.01, and ***p < 0.001 as analyzed by one-way ANOVA.

In recent studies, several proteins that regulate the stabilization, internalization, intracellular trafficking, and recycling of TLR4 have been identified.(Aerbajinai et al., 2013; Cao et al., 2016; Ciesielska et al., 2021; Kim & Kim, 2014; Liu et al., 2023; Tatematsu et al., 2016) We noticed a significant upregulation of CD40, which has been reported to be able to up-regulate membrane expression of TLR4 binding LPS,(Frleta et al., 2003) by IL-2 combined with spike protein compared to spike protein alone in PBMCs (Figure 6A). qPCR analysis confirmed that IL-2 significantly increased the transcription of CD40 in purified monocytes (Figure 6D) and also elevated its surface expression of monocytes in PBMCs (Figure 6E). Importantly, blocking antibodies against CD40 strongly reduced the expression of TLR4 on the membrane surface of monocyte of PBMCs stimulated with IL-2 and spike protein (Figure 6F). The secretion of TLR4 downstream inflammatory cytokines such as IL-6, IL-1β, and IL-8 was also significantly inhibited upon CD40 blocking (Figure S4), underscoring the vital role of CD40 in mediating the TLR4-NF-κB signaling pathway in response to synergistic stimulation of spike protein and IL-2.

Although IL-2 increased the expression of CD40 on monocytes in PBMCs, purified monocytes directly stimulated with IL-2 did not exhibit any significant increase in the expression of CD40 at all (Figure 6G). This finding suggests the involvement of other immune cells in CD40 regulation. Based on the data in Figure 3-5, we hypothesized that TNF-α or IFN-γ might play a role in CD40 regulation. Along this line, we found that TNF-α or IFN-γ stimulated PBMCs markedly upregulated the expression of CD40 on the surface of monocytes in PBMCs, with IFN-γ having a potency similar to IL-2 (Figure 6H). Notably, the expression of CD40 in monocytes stimulated by TNF-α or IFN-γ significantly increased, whereas IL-2 had no effect (Figure 6I, J), suggesting that TNF-α and IFN-γ may mediate the role of IL-2 in upregulating the expression of CD40 in PBMCs, as they are upregulated by IL-2 (Figure 3H). Furthermore, blocking antibodies against TNF-α or IFN-γ effectively attenuated the upregulation of CD40 on the surface of monocyte within PBMCs treated with IL-2 (Fig 6K) and reversed the upregulation of TLR4 on the surface of monocytes in PBMCs treated with spike protein and IL-2 (Figure 6L). These results suggest that the regulation of membrane expression CD40 and TLR4 in response to synergistic stimulation of spike protein and IL-2 is mediated by TNF-α and IFN-γ. Moreover, we found that IKK-16, an inhibitor of NF-κB, could inhibit the promotion of TNF-α and IFN-γ on CD40 expression (Figure 6M).

Considering the vital role of the TLR4-NF-κB signaling pathway in mediating the synergistic stimulation of spike protein and IL-2 in inducing inflammatory cytokines, this pathway presents a promising target for preventing or reducing CRS in COVID-19 patients. Consistent with the effect of IKK-16, we found that TAK242, a TLR4 inhibitor, significantly inhibited the release of IL-1β, IL-6, and IL-8 in PBMCs co-treated with IL-2 and spike protein (Figure 6N). Notably, this inhibitory effect was further enhanced when combined with blocking antibodies against TNF-α or IFN-γ (Figure 6N).

Our findings suggested that IL-2 stimulation of PBMCs leads to a significant secretion of IFN-γ and TNF-α, resulting in the upregulation of CD40 on the surface of monocytes. The increased interaction of CD40-CD40L then maintains the stable membrane localization of TLR4 and prolongs its interaction with spike protein, subsequently hyper-activating NF-κB in monocytes. These findings shed light on the mechanism underlying the synergistic effect of spike protein and IL-2 in inducing inflammatory cytokine release. They may provide important insights for developing effective therapeutic strategies to prevent or reduce CRS in patients with COVID-19.

## Discussion

CRS is a common complication observed in severe COVID-19 patients.(Yongzhi, 2021) IL-6 and IL-1 are considered to be essential cytokines in CRS. We observed that spike protein can stimulate PBMCs to produce IL-1β and IL-6 in a dose-dependent manner. We initially speculated that an increase in viral load could lead to elevated spike protein levels and trigger CRS. However, there was no significant difference in viral load between severe and mild COVID-19 patients (To et al., 2020). Moreover, Clinical reports have shown that the viral load of patients with COVID-19 is relatively high in the initial stage of infection.(Pan et al., 2020; Walsh et al., 2020) These findings indicate that factors other than the viral load may play a more significant role in inducing CRS in COVID-19 patients.

One prominent feature of CRS in severe COVID-19 patients is the apparent increase in levels of cytokines. Many severe COVID-19 patients exhibit a noticeable elevation of IL-2 in their plasma.(Akbari et al., 2020; Huang et al., 2020; J. Liu et al., 2020) It has been reported that high-dose IL-2 administration causes capillary leak syndrome,(Boyman & Sprent, 2012; van Haelst Pisani et al., 1991) and coincidentally capillary leak syndrome is a severe clinical manifestation of CRS (Case et al., 2020). These studies hint at the crucial role of IL-2 in CRS in severe COVID-19 patients. Along this line, our study revealed that DCs stimulated by spike protein activate T cells to release more IL-2 than those stimulated with DCs alone. Therefore, we speculate that spike protein may promote DCs to activate T cells to release a large amount of IL-2, leading to capillary leak syndrome in severe COVID-19 patients. Moreover, capillary leak syndrome also occurs in some individuals who received the SARS-CoV-2 RNA vaccine which translates and expresses spike protein *in vivo*,(Matheny et al., 2021) further supporting our speculation. IL-2 plays a critical role in the immune system, and we found that IL-2 alone can stimulate PBMCs to produce multiple cytokines (IFN-α, IFN-β, TNF-α), which may lead to the persist inflammation. Additionally, IL-2 and spike protein of SARS-CoV-2 synergistically stimulated PBMCs to produce a large amount of IL-1β, IL-6, and IL-8. These results further reveal that the continuous presence of virus antigens and persistent inflammatory reactions are the primary causes of CRS, which is consistent with the clinical symptoms of severe COVID-19 cases. Clinically, severe COVID-19 patients exhibit a slight decrease in viral load but persistent inflammatory reactions.(Moss, 2022; Zheng et al., 2020) Thus, our findings indicate that IL-2 is essential in CRS in patients with severe COVID-19.

Severe COVID-19 patients with CRS often exhibit activation of NF-κB, as well as upregulation of TNF-α and IFN-γ.(Hariharan et al., 2021; Huang et al., 2020; Kircheis et al., 2020) However, the underlying mechanism is vague. In this study, we found that IL-2 activated the NF-κB pathway of monocytes by stimulating PBMCs to release TNF-α and IFN-γ. It has been reported that spike protein of SARS-CoV-2 activates monocytes through binding to TLR4 which is an important upstream regulator of NF-κB.(Brandao et al., 2021; Conte, 2021; Manik & Singh, 2022; Zhao et al., 2021b) We confirmed that spike protein activated the NF-κB, but when combined with IL-2, the NF-κB pathway becomes even more strongly activated. NF-κB inhibitor not only represses the NF-κB activation induced by spike protein or IL-2 combined with spike protein, but also reduces the release of IL-1β, IL-6, and IL-8 from PBMCs stimulated by IL-2 combined with spike protein. These indicate that the NF-κB is vital in the interplay between IL-2 and spike protein in inducing CRS-related inflammatory factors in PBMCs. Targeting the NF-κB pathway could, therefore, prove helpful in preventing and treating CRS in patients with COVID-19. It has been reported that aspirin may reduce mortality in patients with COVID-19,(Martha et al., 2021) but the mechanism is unclear. Since aspirin can inhibit NF-κB activity, (Huo et al., 2018; Liao et al., 2015) we speculate that aspirin may reduce the risk of severe CRS in COVID-19 patients by inhibiting the activity of NF-κB. Overall, our study clarified the interplay between IL-2, TNF-α, IFN-γ, and CRS in COVID-19 patients and highlighted potential avenues for therapeutic intervention.

We found that spike protein of SARS-CoV-2 reduces the membrane expression of TLR4 on monocytes, which may be due to the internalization of TLR4 after binding to its ligand.(Kagan et al., 2008) The internalized TLR4 locates at the endosome and involves in TRIF-dependent signaling pathways, leading to a decreased and delayed activation of NF-κB compared to the TLR4-MyD88 signaling axis.(Lin et al., 2021) In addition, the internalized TLR4 would eventually be degraded by lysosomes, further weakening the NF-κB signaling activated by membrane TLR4.(Gangloff, 2012) Of note, after adding IL-2, the decrease in TLR4 membrane expression caused by spike protein was reversed, and the TLR4-NF-κB pathway was further activated. This occurred because IL-2 maintained the stable localization of TLR4 on the surface of monocytes treated with spike protein by up-regulating the expression of CD40. The interaction between CD40 and CD40L helped stabilize TLR4 on the surface of monocytes treated with spike protein, thus avoiding its rapid internalization and degradation. Our research is consistent with previous reports in which they believe that the binding of CD40 and CD40L can improve the stability of TLR4 on the surface of DCs.(Frleta et al., 2003) Although IL-2 could not directly stimulate the expression of CD40 on monocytes, it achieved this effect through IFN-γ and TNF-α. It should be noted that the impact of TNF-α was weaker than that of IFN-γ, consistent with literature reports.(Lee et al., 2007) These results indicate that TNF-α and IFN-γ are crucial in enhancing the TLR4-NF-κB pathway via transcriptional regulation of CD40. Our study further showed that TNF-α neutralizing antibody combined with a TLR4 inhibitor inhibits the synergistic stimulation of IL-2 and spike protein in monocytes releasing CRS-related inflammatory factors such as IL-1β, IL-6, and IL-8. Therefore, the combination of TLR4 inhibitor and TNF-α neutralizing antibody may represent a promising therapeutic strategy in treating CRS in patients of COVID-19 and other syndromes, such as CAR-T cell anti-tumor therapy, where CRS is a common side effect.

Several clinical studies have shown that monocytes are involved in the developing of CRS in severe COVID-19 patients,(Falck-Jones et al., 2022; Ma et al., 2022; Merad & Martin, 2020; Pence, 2020) but currently, no effective *in vitro* model exists for verification. We found that when monocytes were removed from PBMCs, the synergistic effect of IL-2 and spike protein in stimulating PBMCs to release CRS-related cytokines significantly decreased. This indicated that our model could effectively certify that monocytes are the critical cells for releasing CRS-related cytokines. However, direct stimulation of monocytes with IL-2 combined with spike protein did not enhance the release of inflammatory factors, suggesting that IL-2 is not the primary factor directly cooperating with spike protein to stimulate monocytes to release inflammatory factors. Our study also found that PBMCs stimulated with IL-2 exhibited increased secretion of TNF-α and IFN-γ from NK cells and IFN-γ from T cells. Co-culturing NK cells or T cells with monocytes in the medium containing IL-2 and spike protein significantly increased the release of inflammatory factors, including IL-1β, IL-6, and IL-8. These findings further suggest that NK cells, T cells, and monocytes play an essential role in CRS induced by SARS-CoV-2. Our research provides novel insights into the interplay between immune cells and cytokines produced by CRS, laying the foundation for a better understanding of this severe complication related to COVID-19.

## Conclusion

Our research suggests that after SARS-CoV-2 enters the human body, spike protein can activate DCs, which in turn stimulate T cells producing a large amount of IL-2. IL-2 can stimulate NK cells to release TNF-α and IFN-γ and T cells to release IFN-γ. The increased level of IFN-γ enhances the membrane-located TLR4 via up-regulating the transcription of its partner CD40 in monocytes, resulting in prolonged interaction between spike protein and TLR4 and subsequent activation of NF-κB. Simultaneously, the elevated level of TNF-α activates the NF-κB signaling pathway of monocytes. In such a condition, TNF-α and IFN-γ cooperate to enhance the activation of NF-κB-dependent transcription of CRS-related inflammatory cytokines such as IL-1β, IL-6, and IL-8 (Figure 7). Overall, our study shows that the development of CRS requires the involvement of multiple immune cells, and IL-2, TNF-α, and IFN-γ may act as the primary factors in triggering CRS, providing promising avenues for clinical interventions for COVID-19 patients.

**Figure 7.**
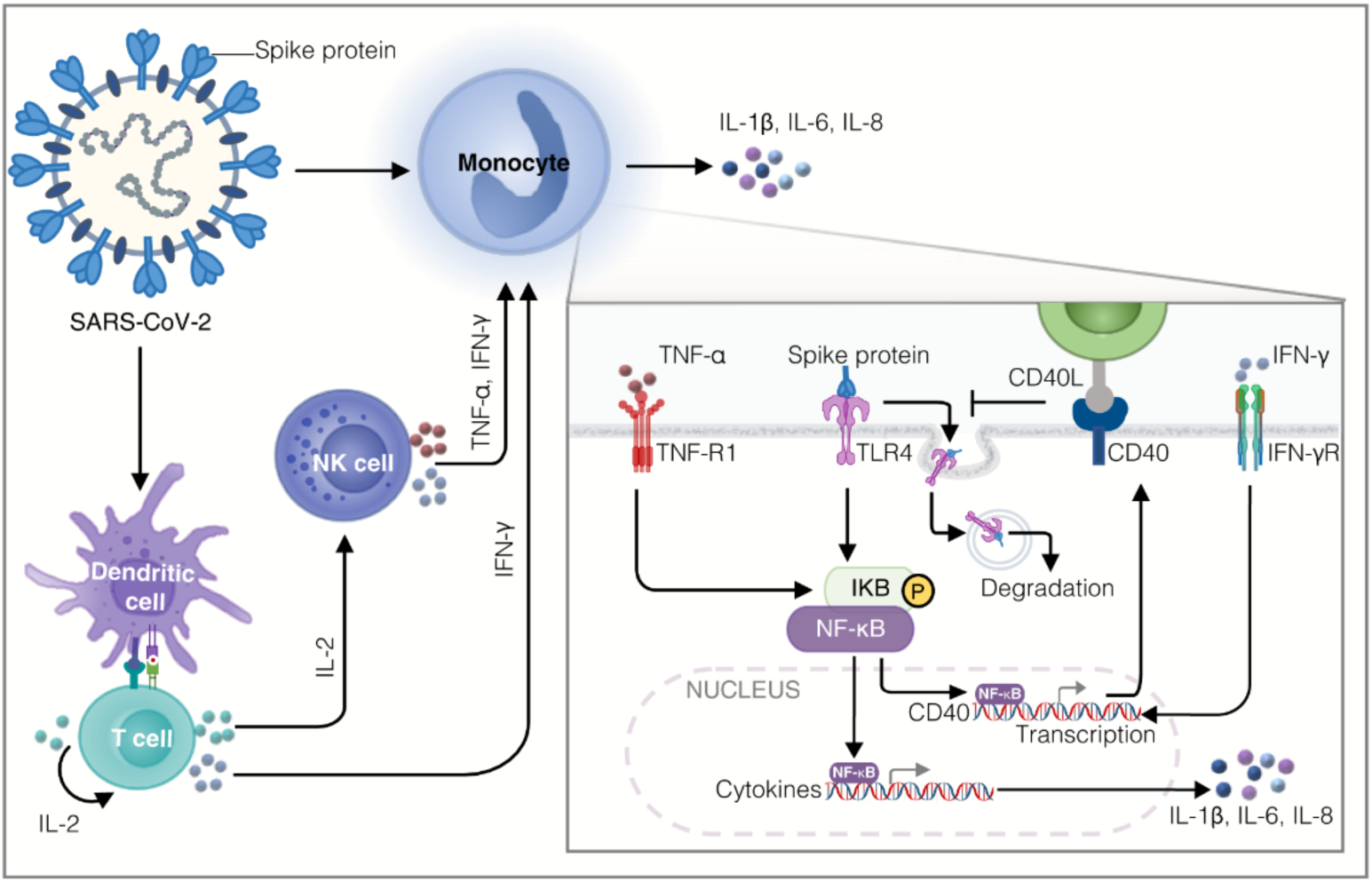
A diagram of the mechanism of spike protein cooperating with IL-2 to stimulate immune cells to produce CRS-related inflammatory factors. DCs loaded with spike protein stimulate T cells to secrete IL-2, which subsequently facilitates the production of TNF-α and IFN-γ by NK cells and IFN-γ by T cells. IFN-γ increases the transcription of CD40, which promotes the stable localization of TLR4 on the membrane surface of monocytes, leading to a constant interaction between spike protein and TLR4 and activation of NF-κB. TNF-α also activates NF-κB signaling in monocytes, which cooperates with IFN-γ to modulate NF-κB-dependent transcription of CRS-related inflammatory cytokines such as IL-1β, IL-6, and IL-8.

## Materials and Methods

### Isolation of PBMCs

The Ethics Committee of the First Hospital of Jilin University approved this study (2020-521). The peripheral blood samples rich in white blood cells were initially obtained from the Changchun Central Blood Station. PBMCs were obtained by Ficoll density gradient centrifugation of lymphocyte separation solution Lymphoprep^TM^ (Axis-Shield, Oslo, Norway). PBMCs were cultured in RPMI-1640 medium (Gibco, Grand Island, NY, USA) in a humidified atmosphere with 5% CO2 at 37°C.

### Spike protein stimulates PBMCs

PBMCs were adjusted to 1×10^6^ cells/mL RPMI-1640 medium, and the PBMCs were inoculated into 24-well plates (NEST Biotechnology Co., Ltd, Wuxi, China) at 400 μL per well. Then PBS, spike protein (ACROBiosystems, Beijing, China), IL-2 (100 IU/mL), IL-2 (100 IU/mL) combined with spike protein (10 μM), TNF-α (0.5 ng/mL), TNF-α (0.5 ng/mL) combined with spike protein (10 μM), IFN-γ (1 ng/mL), or IFN-γ (1 ng/mL) combined with spike protein (10 μM) (above cytokines were all from T&L Biological Technology, Beijing, China) were added to PBMCs. After 16 hrs of incubation, the supernatant was collected for cytokine detection. In some experiments, blocking antibodies against CD40 (10 μg/mL) (Biolegend, San Diego, USA), TNF-α (10 μg/mL) (MedChemExpress, Monmouth Junction, USA), or IFN-γ (10 μg/mL) (eBioscience, San Diego, USA) were added to the cultures.

### Isolation of NK cells, T cells, and monocytes

T cells, NK cells, and monocytes were isolated from PBMCs by EasySep Human T Cell Isolation Kit (StemCell Technologies, Vancouver, Canada), MACSxpress Whole Blood NK Cell Isolation Kit (Miltenyi Biotec, Bergisch Gladbach, Germany), and EasySep Human Monocyte Isolation Kit (StemCell Technologies). In the coculture experiment, the density of monocytes was adjusted to 2×10^5^ cells/mL, and NK or T cells were mixed with monocytes of corresponding donors according to the ratio of NK or T cells to monocytes. Cells were inoculated in 24-well plates at 400 μL per well. Then PBS, spike protein, IL-2, or IL-2 combined with spike protein were added respectively and incubated for 16 hrs. The supernatant was collected, and the cytokines in the supernatant were detected.

### DCs and T cell coculture

DCs were adjusted to 1×10^6^ cells/mL in CellGenix® GMP DC serum-free medium (CellGenix, Freiburg, Germany) with 100 ng/mL GM-CSF and 50 ng/mL IL-4 (both from T&L Biological Technology). Spike protein and TNF-α were added on the fifth day of culture, and DCs were collected on the seventh day. The density of T cells was adjusted to 1×10^6^ cells /mL, and DC and T cells were cocultured at a ratio of 1:5 for two days. The supernatant was collected for the detection of IL-2.

### Flow cytometry

The cell density should be adjusted to 1×10^6^ cells/mL, and then 1 μL BD GolgiPlug™ (BD Biosciences, San Jose, CA, USA) should be added to each milliliter of the cell suspension. Cells were cultured in an incubator at 37℃ for 4 h and washed with PBS once. Following the instructions of BD Cytofix/Cytoperm™ Plus Fixation/Permeabilization Solution Kit (BD Biosciences), Cells were fixed, permeabilized, and stained with anti-human TNF-α (Biolegend), anti-human IFN-γ (BD Biosciences), anti-human IL-6 (Biolegend), or isotype control antibodies for 30 min.

To detect CD40 and TLR4 on monocytes, cells were adjusted to 1×10^6^ cells/mL and stained with anti-human CD14, anti-human CD40, anti-human TLR4, or isotype control antibodies (the above antibodies were all from Biolegend) at room temperature for 15 min.

PBMC should be adjusted to a cell density of 2.5×10^6^ cells/mL. An equal volume of BD Cytofix™ Fixation Buffer (BD Biosciences) was added to the cell suspension and fixed at 37°C for 10 min. After fixation, the cells were collected and mixed with 200 μL BD PhosFlow^TM^ Perm Buffer III (BD Biosciences), then incubated on ice for 30 minutes. The cells were washed with PBS twice and incubated with anti-human NF-κB p65 (pS529) (BD Biosciences), anti-human CD14, or isotype control antibodies for 30 min at room temperature in the dark.

After washing with PBS once, the cells were analyzed using a FACSAria II flow cytometer (BD Biosciences). The data were analyzed by FlowJo software version 10 (Tree Star, Inc., Ashland, OR, USA).

### Cytokine detection

Cytokines in the supernatants were assayed by Cytometric Bead Array (CBA, BD Biosciences) or respective ELISA kit (Biolegend) according to the manufacturer’s instructions.

### Quantitative Real-time PCR

PBS, spike protein, IL-2, or IL-2 combined with spike protein were used to stimulate PBMCs for 16h. Then monocytes were isolated by a monocyte enrichment kit. RNA of monocytes was extracted by GeneJET RNA Purification Kit (Thermo Fisher Scientific, Waltham, MA, USA), and cDNA was obtained by Hifair 1st Strand cDNA Synthesis SuperMix for qPCR (Yeasen, Shanghai, China). Quantitative real-time PCR (qPCR) was performed using 2×RealStar Green Fast Mixture (GenStar, Beijing, China) in a CFX384 Real-Time System C1000 Touch Thermal Cycler (Bio-Rad Laboratories, Hercules, CA, USA). The relative transcription levels of the genes of interest were normalized against the expression of GAPDH and calculated using the ΔΔCT method. The sequences of primers used in this study are CD40, CCTGTTTGCCATCCTCTTGGTG (forward primer) and AGCAGTGTTGGAGCCAGGAAGA (reverse primer); TLR4, AGACCTGTCCCTGAACCCTAT (forward primer) and CGATGGACTTCTAAACCAGCCA (reverse primer); GAPDH, GTCTCCTCTGACTTCAACAGCG (forward primer) and ACCACCCTGTTGCTGTAGCCAA (reverse primer).

### High throughput sequencing

PBMCs were stimulated with PBS, spike protein, IL-2, or IL-2 combined with spike protein for 16 hrs and used for RNA extraction using TRIzol™ reagent (Thermo Fisher Scientific, MA, USA) according to the manufacturer’s instructions. RNA sequencing and bioinformatic analysis were performed at Shanghai Majorbio Bio-Pharm Technology Co., Ltd. (Shanghai, China). RNA sequencing data has been uploaded to the GEO database. GEO accession numbers is GSE241843.

### Statistical analyses

Statistical analysis was performed by GraphPad Prim 8 (GraphPad Software). Paired Student’s t-test was performed for the comparison of two paired groups. One-way ANOVA or two-way ANOVA was conducted to compare three or more groups. *P* value of less than 0.05 was considered statistically significant.

## Acknowledgments

This work was supported by the National Key Research and Development Program of China under Grant number 2020YFA0707704, the Innovative Program of National Natural Science Foundation of China under Grant number 82050003, National Natural Science Foundation of China under Grant numbers 82273186, 82002962, and 81874052, Project funded by China Postdoctoral Science Foundation under Grant number 2021M691208, Jilin Provincial Science and Technology Department under Grant numbers 20200602032ZP, 20210401076YY, 20210303002SF and YDZJ202202CXJD004, Jilin Provincial Education Department under Grant number JJKH20221060KJ, Jilin Province Labor Resources and Social Security Department under Grant number 2023RY03, Jilin Provincial Development and Reform Commission under Grant number 2021C010, and Changchun Science and Technology Bureau under Grant number 21ZGY28.

## Data Availability Statement

All data generated or analyzed during this study are included in this published article.

## Figure Legends

**Figure S1.**
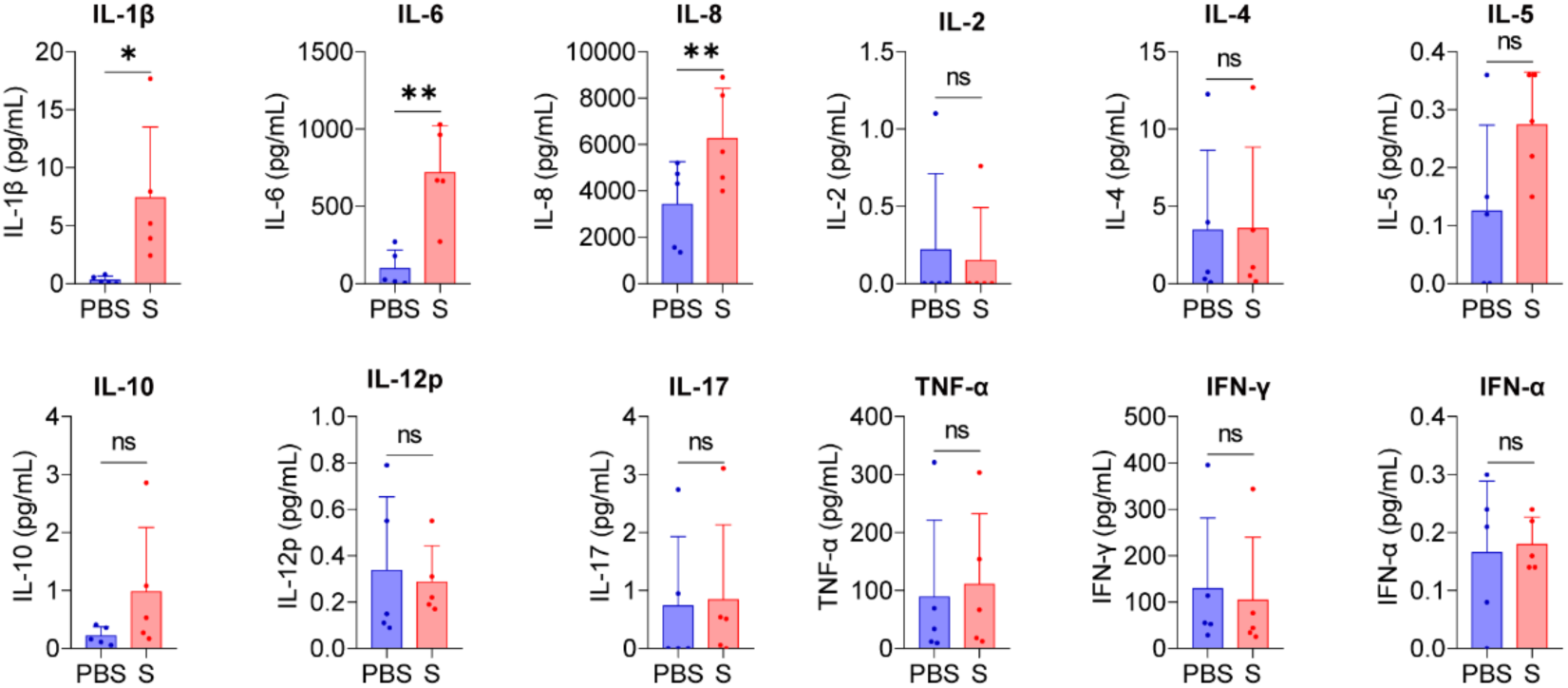
Spike protein stimulates PBMCs to secrete IL-1β, IL-6, and IL-8. Quantifying concentrations of cytokines by CBA in the supernatants of PBMCs stimulated by spike protein for 16 hrs (n = 5 biological replicates). Data are presented as mean ± SD. ns: not significant, *p < 0.05, and **p < 0.01 as analyzed by paired Student’s t-test.

**Figure S2.**
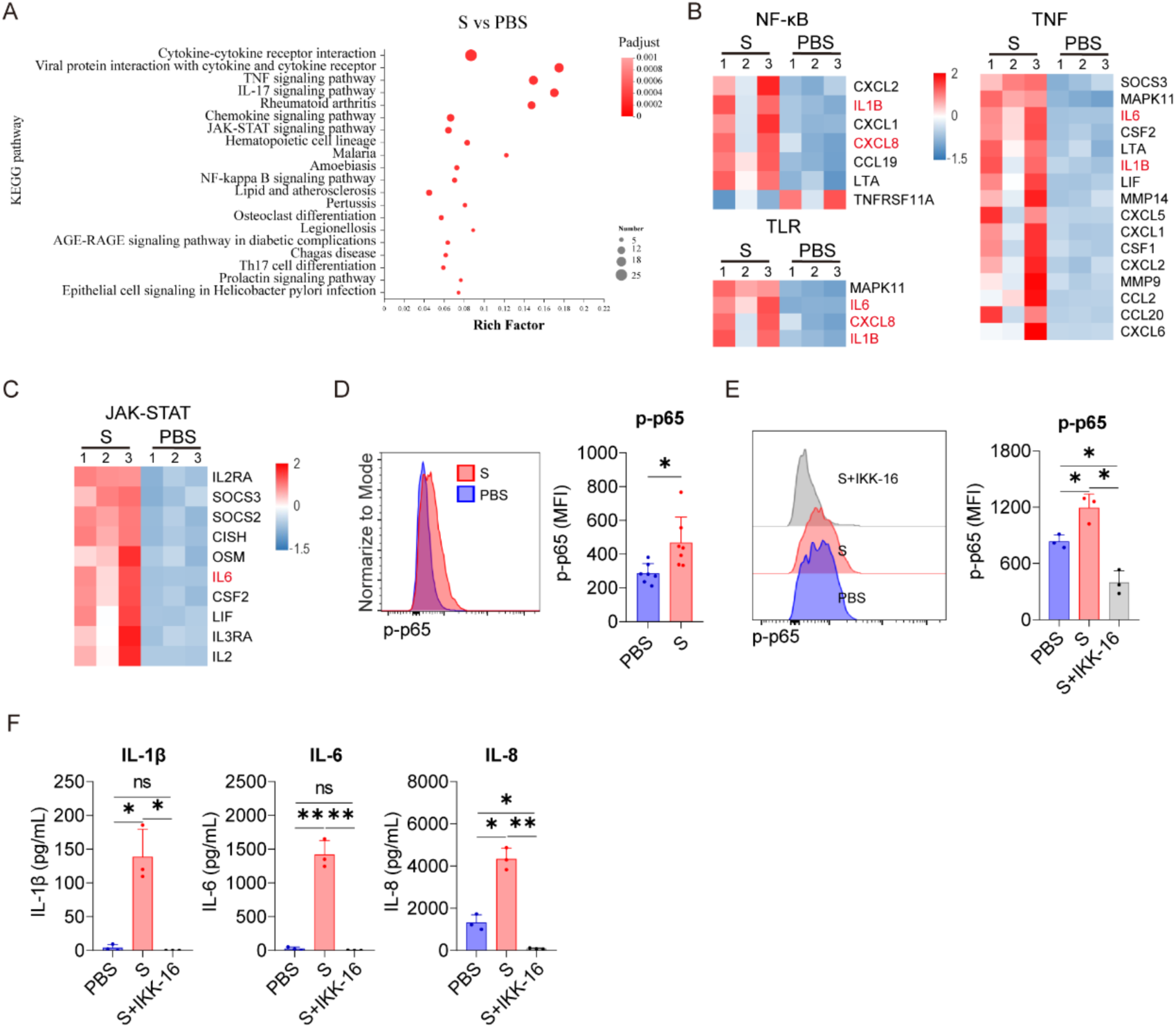
Spike protein activates NF-κB to facilitate monocyte transcription of IL-1β, IL-6, and IL-8. (A) Bubble plot showing the enrichment of KEGG pathways based on the transcriptomic analysis of PBMCs treated with spike protein or PBS for 16 hrs. (B, C) Heat maps showing the differentially expressed genes in the NF-κB signaling pathway, TNF-α signaling pathway, and JAK-STAT signaling pathway in PBMCs treated with spike protein or PBS for 16 hrs (n = 3 biological replicates). (D) Representative intracellular staining analysis (left) and quantification (right) of the expression of p-p65 by flow cytometry in monocytes of PBMCs stimulated with spike protein or PBS for 16 hrs (n = 7 biological replicates). (E) Representative intracellular staining analysis (left) and quantification (right) of the expression of p-p65 by flow cytometry in monocytes of PBMCs stimulated with PBS, spike protein, or spike protein combined with IKK-16 (n = 3 biological replicates). (F) Quantifying concentrations of IL-1β, IL-6, and IL-8 by CBA in the supernatants of PBMCs stimulated with PBS, spike protein, or spike protein combined with IKK-16 for 16 hrs (n = 3 biological replicates). Data are presented as mean ± SD. ns: not significant, *p < 0.05, **p < 0.01, and ***p < 0.001 as analyzed by one-way ANOVA (E, F) or paired Student’s t-test (D).

**Figure S3.**
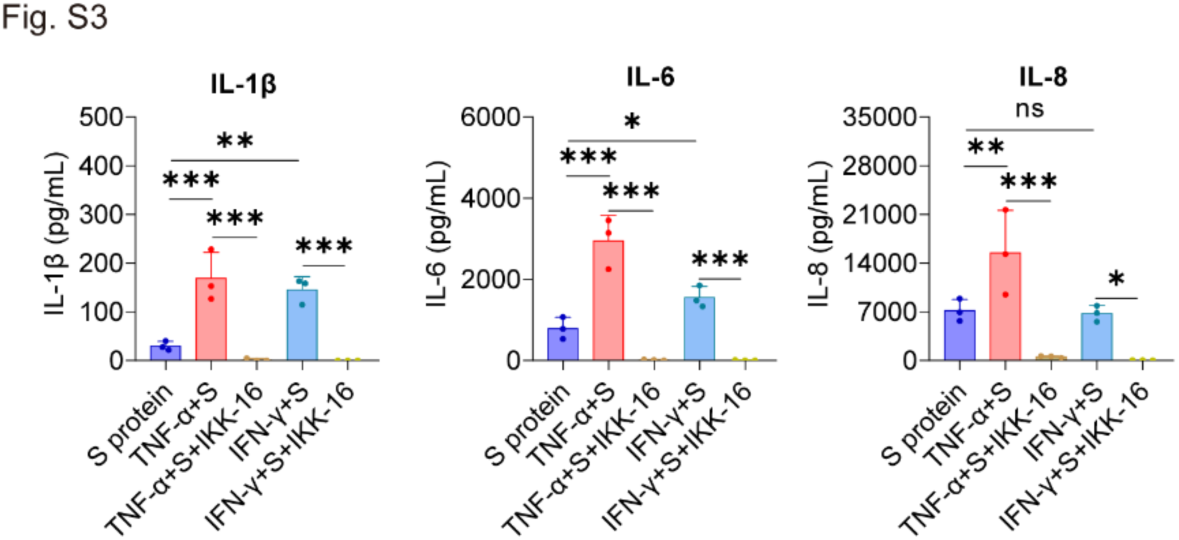
Inhibition of NF-κB reduces the secretion of IL-1β, IL-6, and IL-8 in PBMCs stimulated by spike protein together with TNF-α or IFN-γ. Quantifying concentrations of IL-1β, IL-6, and IL-8 by CBA in the supernatants of PBMCs stimulated with spike protein, spike protein combined with TNF-α or IFN-γ, and spike protein combined with TNF-α or IFN-γ together with IKK-16 for 16 hrs (n = 3 biological replicates). Data are presented as mean ± SD. ns: not significant, *p < 0.05, **p < 0.01, and ***p < 0.001 as analyzed by one-way ANOVA.

**Figure S4.**
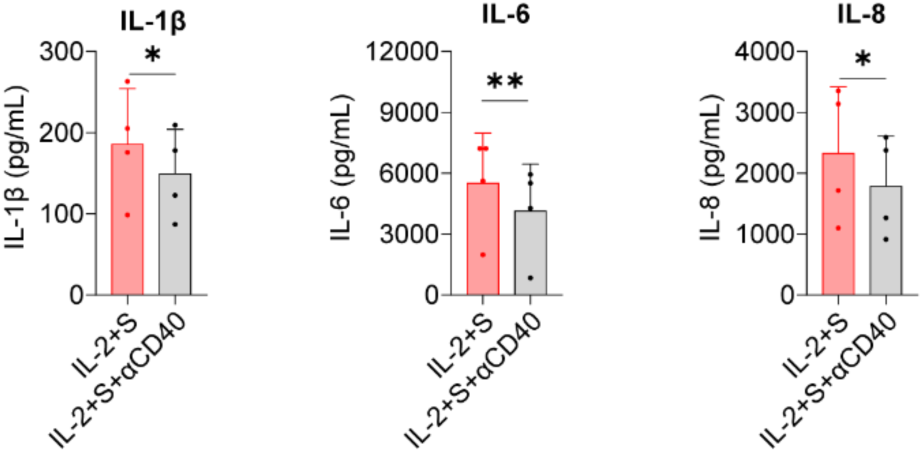
Reduction in the secretion of IL-1β, IL-6, and IL-8 in PBMCs stimulated with IL-2 and spike protein upon blocking CD40. Quantifying concentrations of IL-1β, IL-6, and IL-8 by CBA in the supernatants of PBMCs stimulated with spike protein combined with IL-2 with/without CD40 blocking antibody for 16 hrs (n = 4 biological replicates). Data are presented as mean ± SD. *p < 0.05, and **p < 0.01 as analyzed by paired Student’s t-test.

## KEY RESOURCES TABLE

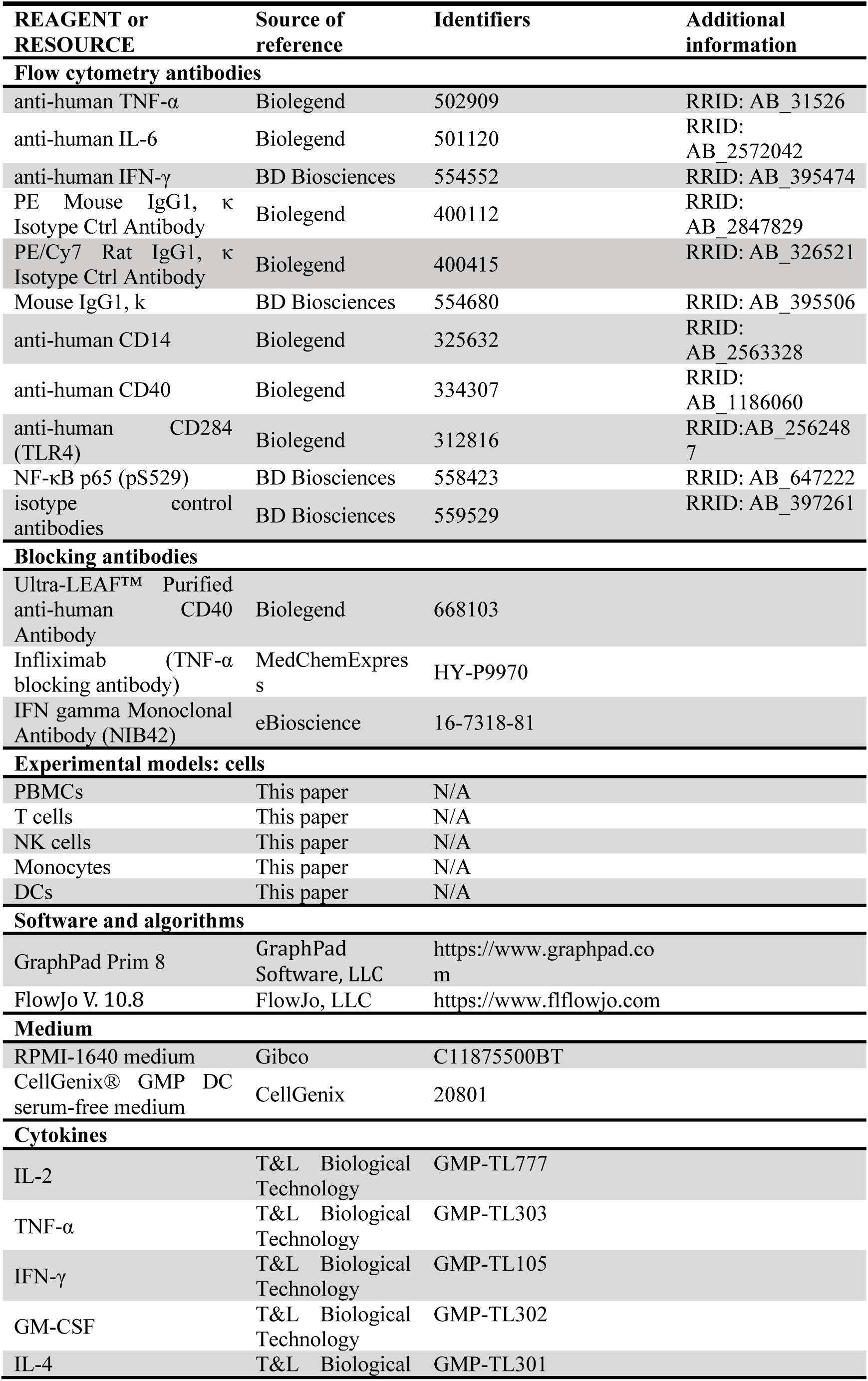
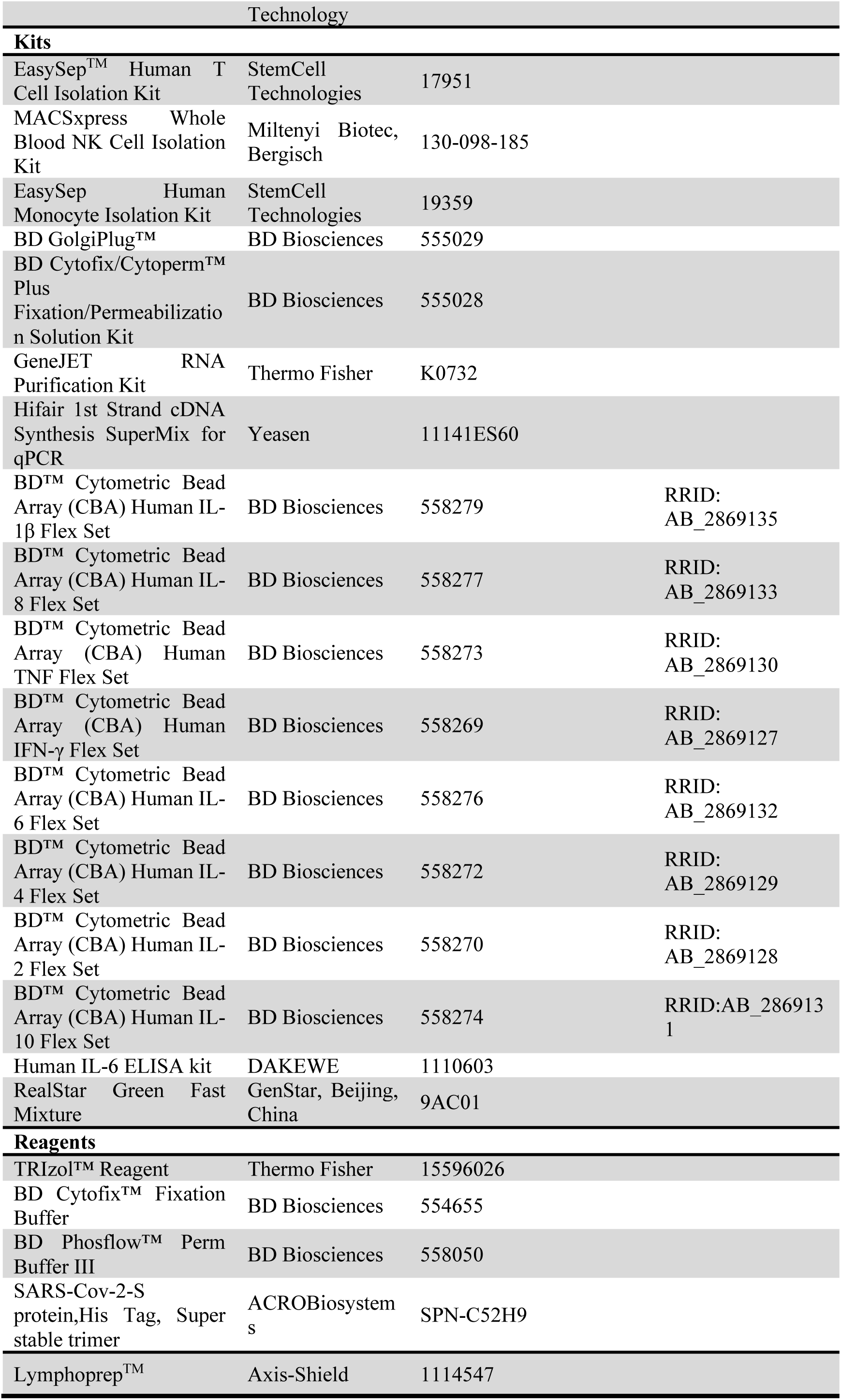

